# The Human Bindome: A Proteome-scale Atlas of Designed Binder Candidates

**DOI:** 10.64898/2026.07.30.741542

**Authors:** Julius Wenckstern, Anna M. Díaz-Rovira, Julia Kuhn, Arvid Ban, Rahma Hamdani, Roser Pruaño-Milla, Evgenia Elizarova, Sandrine Georgeon, Kaiden Thompson, Matthias Hinterndorfer, Devanarayanan Siva Sankar, Maximilian Dünnebacke, David Desscan, Sreenath Nair, Marcelo Querino Lima Afonso, Jennifer Fleming, Sameer Velankar, Andrea Ablasser, Paola Picotti, Georg Winter, Mikko Taipale, Bruno E. Correia

**Author notes:** These authors contributed equally (Julius Wenckstern, Anna M. Díaz-Rovira).

## Abstract

Affinity reagents such as antibodies are indispensable for interrogating proteins’ biological function. Yet they are costly and frequently unreliable, with unknown sequences, posing challenges to reproducible experimental research. Deep learning-based protein design can now *in silico* generate affinity reagents achieving reliable experimental success rates, but has remained largely confined to specialist laboratories. Here we present the Human Bindome, a proteome-scale atlas of high-confidence *in silico* protein binder candidates. By embedding the experimentally benchmarked BindCraft method in an accelerated, parallelized framework with automated domain-level target selection, we generated 306,146 binder candidates covering 8,296 human proteins (40.9% of the full proteome). Every candidate carries a defined sequence, a predicted binder-target structure model, and *in silico* confidence metrics. We characterize proteome-wide coverage and show that binder epitopes frequently overlap functional sites. This positions the Bindome as a resource of genetically encodable perturbagens for site-specific, modular control of protein function. The Bindome is freely available through a web interface (https://bindome.epfl.ch), with agentic, natural-language querying and as data splits for machine-learning model development. We anticipate that the Bindome will be valuable for the scientific community by providing affinity and perturbation reagents with broad applications in dissecting biological mechanisms as well as in drug and target discovery.

## INTRODUCTION

Protein-based affinity reagents, such as antibodies and other binders, are powerful tools for investigating and manipulating individual proteins in biological research^1^. These reagents support applications ranging from detection, imaging, and enrichment to functional perturbation, structural stabilization, synthetic biology, and therapeutic development. Despite their versatility, high-quality binders have traditionally remained time, labour, and cost intensive to generate, constraining the number of proteins, epitopes, and distinct species that can be studied.

Many commercially available reagents are poorly characterized^2,3^, with their sequence and molecular composition often unknown^4^. Large repositories exist but each carries trade-offs. SAbDab^5^ catalogues antibody sequences and structures together with their antigens aggregated from the PDB, but it is antibody-centric and covers only about 2.4% of the human proteome (493 target proteins). Antibodypedia^6^ reports around 91.9% coverage (18,640 target proteins), yet its vendor aggregated monoclonal and polyclonal antibodies commonly lack in-depth characterization and sequences while suffering from high batch-to-batch variability. ABCD^7^ ensures sequence availability by aggregating only chemically defined antibodies, but covers just 6.9% (1,409 target proteins). Overall, gaps in characterization, reproducibility, and availability across proteins, epitopes, and species significantly limit our ability to interrogate biological systems^3,8^.

Historically, biology has been studied through perturbation^9,10^. For instance, genetic knockouts^11^, knockins^12,13^, and mutagenesis^14^ have enabled functional studies that disentangle the complexity of biological systems. Genome-wide CRISPR and RNAi approaches demonstrated that omics-scale coverage enables functional screens revealing principles of cellular organization not accessible through individual-target experiments^11,15^, and approaches such as base editor screens^14^ and emerging multiplexed targeted degradation^16^ have further expanded the repertoire of protein-level perturbations. Yet direct and programmable perturbation of a large number of distinct proteins and epitopes remains limited by the availability of binders at proteome scale. Custom-made, site-targeted reagents across proteomes could open many routes to probe biological function in complex systems, enabling protein-level control over interactions, activity, localization, and effector recruitment. Beyond basic discovery, such reagents bear directly on drug and target discovery. Because binders can be directed to defined functional sites, they offer a systematic way to assess the tractability of candidate targets, including disease-associated proteins that lack small-molecule pockets, and to prototype the modulation of protein function that a future therapeutic would need to achieve.

Recent advances in deep-learning-based protein structure prediction and design have begun to remove this bottleneck, transforming binder generation from a specialized, target-by-target screening problem into an on-demand design problem. Methods such as BindCraft^17^, BoltzGen^18^, RFdiffusion^19,20^, and Proteina-Complexa^21^ can generate high-affinity binders directly from target structures, whether experimentally determined or predicted with high confidence. Computational binder design often achieves binding success in more than 10%^17^ of experimentally tested designs. The main obstacles to applying these methods at proteome scale are the technical expertise required and the computational cost of binder generation. A precomputed atlas of binder candidates therefore extends current design tools to proteome scale, making the resulting candidates, and the libraries composed of them, directly accessible. Closest to a precomputed resource, Balbi et al.^22^ mapped ∼4,500 targetable sites across the human surfaceome and provided binder seeds to initiate design, but focused on the cell-surface localized subset and delivered binding seeds rather than complete, ready-to-test binder candidates.

Here we present the Bindome, a comprehensive, *in silico* quality-controlled collection of designed protein binder candidates targeting approximately 77.8% of the structured human proteome. Scaling the previously experimentally validated BindCraft design pipeline, we generated a large collection of *in silico* high-confidence binder candidates spanning diverse structural and functional target classes. We characterized the proteome-wide coverage, analysed the structural features of the binders, and summarized the functional sites that many of them engage. To maximize accessibility, the Bindome is freely available through interactive, programmatic, and agentic interfaces, as well as machine-learning-ready data splits. We anticipate that the Bindome will enable many new detection and perturbation experiments, from single-target studies to pooled screens, with applications spanning basic biology and target discovery, contributing to our understanding of how complex biological systems work and can be modulated.

## RESULTS

### Proteome-scale binder design

To design *in silico* binder candidates at the proteome scale, we developed an accelerated and automated binder design workflow (Fig. 1a). The method builds on the BindCraft generation and filtering framework, which reported experimental success rates ranging from 10-100% across experimentally tested targets^17^, and extends it to proteome-scale execution through automated target selection and preprocessing as well as an accelerated, parallelized implementation. This workflow generated the candidate binders that constitute the Bindome database, in which each binder candidate is linked to a human target protein as identified by UniProt accession or UniProt entry name. Each entry comprises a predicted binder-target-domain complex superimposed onto the full target structure as well as associated *in silico* metrics. All binders were designed at a fixed length of 90 amino acids, ensuring computational efficiency and cost-effective downstream gene synthesis compatible with 300bp oligonucleotide pools.

**Figure 1.**
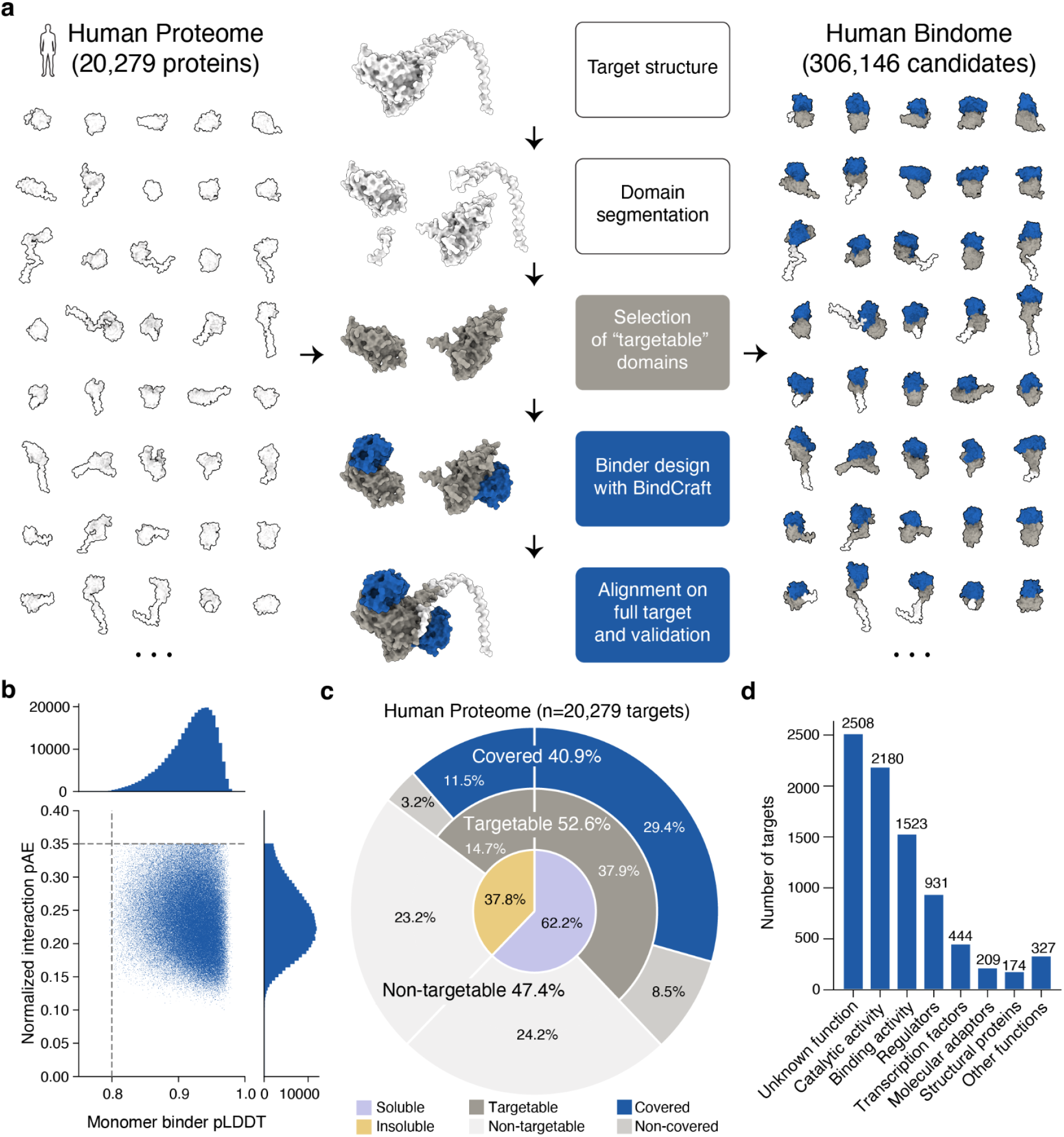
Generation of the Bindome and coverage of the human proteome. **a**, Computational workflow for the design of protein binders at human-proteome scale based on BindCraft. All candidate binders are made available in the Bindome database. **b**, Distribution of the *in silico* confidence metrics across all binder candidates. **c**, Human proteome coverage in the Bindome. Targets were partitioned into soluble and insoluble according to whether the target is membrane-localized. The “targetable” proteome is defined as folded, globular protein domains with confident AlphaFold prediction and comprises all domains for which binder design was attempted. **d**, Functional categories of the targets included in this resource.

Our method uses full-length human protein models and Predicted Aligned Error (PAE) matrices from the AlphaFold (AF) Database^23^ to segment proteins into structural domains by PAE-based clustering^24^. We define targetable domains as compact, confidently predicted regions that pass filters on AF model confidence, domain length, radius of gyration, contact density, secondary-structure composition, and that do not contain membrane-embedded regions. Domains failing these criteria are excluded. This automated preprocessing converts full-length human proteins, including multidomain and partially disordered proteins, into domain-level target models suitable for binder generation using the current state-of-the-art tools.

The domain-level strategy provides two advantages for proteome-scale design. First, it enables binder generation against multiple structurally distinct regions of a target protein without requiring predefined hotspot residues. Second, the reduced target size substantially lowers the computational cost of each design trajectory, which is important because binder-design runtime increases with target length superlinearly. However, designing against isolated domain crops also introduces the possibility that a binder may clash with another region of the full-length protein. To address this, each predicted binder-domain complex is superimposed onto the corresponding full target structure and we reject candidates that clash with structured regions outside the targeted domain. Candidates were retained only when the clashing region was uncertain in its arrangement relative to the interface (high PAE) or was itself of low structural confidence (low pLDDT), such that the apparent clash likely reflects uncertainty in the full-length model rather than a real steric clash (Extended Data Fig. 1).

### *In silico* confidence of the binder candidates

We adopted the BindCraft filters without modification, preserving the stringency of the experimentally benchmarked design framework. Every binder candidate therefore has an AF monomer pLDDT > 0.8 (binder-fold confidence) and a normalized interaction predicted aligned error (iPAE) < 0.35 (binder-target placement confidence) in the complex prediction (Fig. 1b). These distributions were not skewed towards the thresholds (median pLDDT 0.92, median iPAE 0.23) and all remaining BindCraft filtering thresholds were likewise retained (Extended Data Figs. 2, 3). Together, these metrics provide a quantitative basis for prioritizing candidates within each target.

### Proteome-scale target coverage

Thus far, the Bindome contains 306,146 *in silico* binder candidates generated against structurally well-defined, non-membrane-embedded domains selected from soluble and membrane (insoluble) proteins. We define proteins containing at least one such domain as the targetable proteome, comprising 10,664 of 20,279 human target proteins. The resource covers 8,296 target proteins, representing 77.8% of the targetable proteome and 40.9% of the full human proteome (Fig. 1c). It contains an average of approximately 30 candidates per targeted domain and 37 per target protein, yielding a large set of candidates per protein for downstream evaluation and potential applications.

Covered targets span all major molecular function classes of the Gene Ontology^25^ (Fig. 1d). The largest group of targets with designed binders has no assigned function (n = 2,508); among annotated targets, the Bindome covers for instance proteins with catalytic activity (n = 2,180), binding activity (n = 1,523), regulators (n = 931), and transcription factors (n = 444).

We compared the Bindome’s target coverage with established affinity reagent databases. Among these, coverage of the human proteome varies by more than an order of magnitude: Antibodypedia, which aggregates commercial antibody records, lists reagents for 18,640 targets (91.9%), whereas curated or structurally defined collections are far smaller: ABCD (1,409; 6.9%), SAbDab (493; 2.4%), and Biocompare (579; 2.9%). With 8,296 covered targets (40.9%), the Bindome is the second-largest resource by target coverage and by far the largest source of binder candidates with openly available sequences and structural models.

To estimate the structural diversity of the covered targets, we assigned human target domains to CATH superfamilies by structural search against CATH reference domains^26,27^ with Foldseek^28^. Of 1,055 CATH superfamilies represented in the human target-domain set, 78.2% contained at least one domain with a binder candidate, whereas 5.4% were targetable but not yet covered by binders (Fig. 2a). Thus, the Bindome already reaches nearly all targetable structural superfamilies represented in the human proteome. Additionally, we embedded human protein structures in a 2D UMAP based on all-against-all TM-score distances and highlighted targets with at least one binder candidate (Extended Data Fig. 4). Covered targets were distributed throughout the embedding, indicating broad representation across human structural space, with untargeted regions corresponding mainly to targets outside the targetable scope, such as integral membrane proteins.

**Figure 2.**
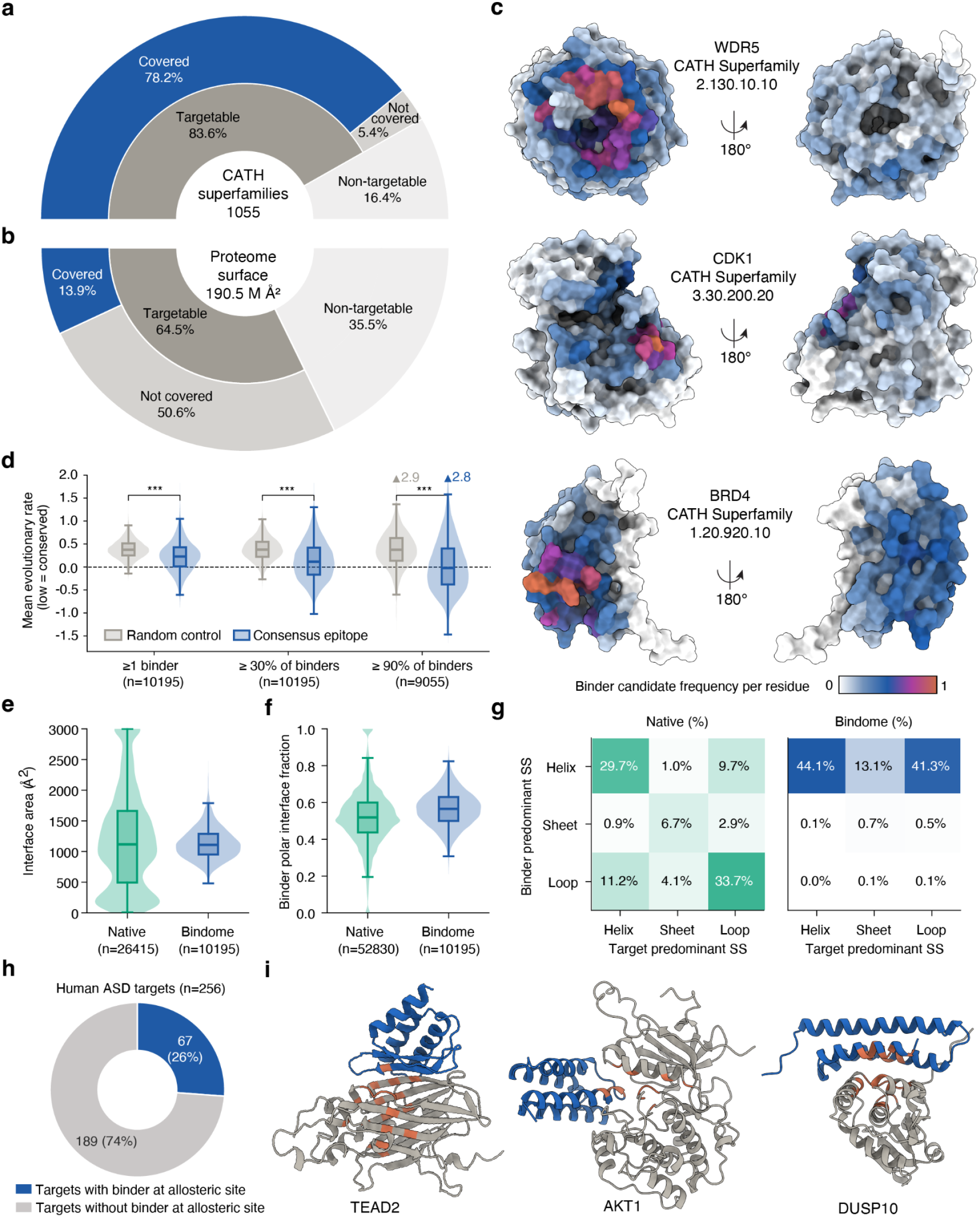
Structural coverage, binder and interface analysis. **a**, Coverage of CATH superfamilies present in the human target-domain set; covered denotes a superfamily containing ≥1 domain with a binder candidate. **b**, Coverage of proteome surface area computed from all full-length AlphaFold models. The inner ring splits the total solvent-accessible proteome surface into targetable and non-targetable under the current design scope; the outer ring splits the targetable surface into the fraction covered by at least one designed binder versus not covered. **c**, Per-residue binder-contact frequency projected onto representative structures for three CATH superfamilies: WDR5 (WD40), CDK1 (protein kinase), and BRD4 (bromodomain). Contacts from binder-target complexes across members of each superfamily were aggregated and mapped onto a representative structure of the domain. **d,** Mean evolutionary rate of binder epitopes compared with residue-count-matched surface control (higher rate = more variable, lower rate = more conserved). Because most target domains have multiple binders, we defined the epitope at three consensus thresholds: residues contacted by at least one binder, by ≥30% of a target’s binders, and by ≥90% of a target’s binders. Two-sided Wilcoxon signed-rank test, paired by domain; all P < 1×10^⁻300^ (below the double-precision floating-point floor). ***P < 0.001. **e**, Interface area for Bindome binder candidates compared with native protein-protein interactions. **f**, Fraction of polar residues at the interface for native protein-protein interactions compared with Bindome binder candidates. **g**, Pairing of secondary structures at the interface. Each interface is classified by the predominant secondary structure of the binder and target. **h,** Coverage of targets in ASD with binders overlapping annotated allosteric-sites. **i,** Representative examples showing target (grey), binder (blue), and residues within 5Å of a known allosteric modulator from a PDB structure (orange): AKT1 (PDB 6HHH), TEAD2 (PDB 6E5G), and DUSP10 (PDB 6MC1).

We quantified how much of the human proteome surface is contacted by the Bindome’s binder candidates. Using full-length AlphaFold models as the reference, we found that 64.5% of the modelled surface is targetable under the current design scope, yet binder contacts covered only 13.9% of the total surface area (Fig. 2b). This is relevant because the utility of a binder resource depends not only on the number of proteins targeted, but also on the diversity of epitopes available on each protein. Each epitope provides an independent handle for perturbing, detecting or recruiting a target protein; accordingly, broader surface coverage increases the number of ways in which the proteome can be interrogated directly at the protein level. Thus, although the Bindome already spans many human proteins, per target epitope coverage remains sparse.

### Epitope distribution and conservation

To better understand what characterizes targeted epitopes, we projected epitope residues from binder-target complexes involving different members of the same CATH superfamily onto representative structures for WD40 domains, protein kinases, and bromodomains. These projections revealed binding hotspot patterns within each family (Fig. 2c). For instance, in bromodomains and WD40 domains, targeted epitopes focused on exposed grooves and pockets that are recurrently used for molecular recognition. Kinase-domain projections showed pronounced epitope consensus across exposed regions of the catalytic domain, including surfaces outside the active-site cleft.

Next, we characterized evolutionary conservation of the epitopes (Fig. 2d). Because each target domain is typically engaged by many binders whose epitopes often overlap, we defined the epitope at three levels of increasing consensus: residues contacted by any binder, by at least 30% of a target’s binders, and by at least 90% of a target’s binders. At every threshold, epitope residues were on average significantly more conserved than a residue-count-matched randomly drawn control of the same protein surface (two-sided Wilcoxon signed-rank test paired by domain, all P < 1×10^-300^). Moreover, conservation increased with consensus: the more binders converged on a set of residues, the more conserved it tended to be. Together, this indicates that designed binders preferentially engage conserved, and therefore likely functionally important, surface regions.

### Designed versus natural protein-protein interactions

To assess whether designed binder-target interactions resemble those in nature, we compared the Bindome interfaces with native protein-protein interactions from PINDER^29^. The Bindome interfaces showed a narrower interface-area distribution, likely owing to the manually fixed 90-residue binder size, though it was centred at the native mean (Fig. 2e). The fraction of polar residues at the Bindome interfaces was also more narrowly distributed, with a modestly higher mean (Fig. 2f).

Whilst native interactions drew on a diverse repertoire of secondary-structure pairings, the Bindome interfaces were strongly enriched for helical binder segments, most often contacting helical or loop regions on the target (Fig. 2g). Consistent with this, binder folds largely matched existing folds, with only 17.3% of binders having a max query TM-score < 0.5 against the AF Database, yet the resulting interfaces were largely novel with a median interface TM-score around 0.3 against the PPIRef^30^ (Extended Data Fig. 5). Altogether, the Bindome interfaces are native-like in size and polarity but show a strong bias on the binder side towards helix-mediated contacts.

Finally, as induced fit is common in native protein interactions, we asked whether the Bindome’s complexes show conformational adaptation (Extended Data Fig. 6). Using a threshold of interface Cα RMSD > 2 Å between bound and unbound states in the AF models, 4% of all binder candidates and 8% of all targets showed conformational adaptation. Target-side adaptation is particularly notable because the target is templated in the complex structure prediction, a bias that would tend to suppress larger conformational adaptations.

### Coverage of allosteric sites

The functional relevance of an epitope depends on where it lies on the target. We therefore asked whether binder epitopes coincide with allosteric sites. Among human targets with an annotated allosteric site in the Allosteric Database (ASD)^31^, 26% of targets carry at least one binder candidate overlapping with the allosteric site (Fig. 2h). Consistent with this, binder candidate epitopes are placed directly over residues that contact a known allosteric modulator, as illustrated for AKT1, TEAD2, and DUSP10 (Fig. 2i). These cases show that binder candidates can engage surfaces of established regulatory importance.

### Functional landscape of candidate binders for perturbation studies

In biological research, perturbations such as mutational and gene-knockout studies have transformed our understanding of fundamental processes in cell biology. The scale of the Bindome extends this potential beyond individual target studies. Because each designed binder candidate engages a defined target region, binder candidates can be treated as putative perturbagens that may generate biological hypotheses at the target-level as well as site-specific protein function. To illustrate the scope of these hypotheses, we integrated the Bindome’s target structures with annotations from external resources including UniProt^32^, Pharos^33^, Mondo^34^, PINDER^29^, PDB^35^, and DepMap^36^. We stratified annotations according to whether they describe properties of the target protein as a whole or features of the target sites engaged by a binder candidate (Fig. 3a).

**Figure 3.**
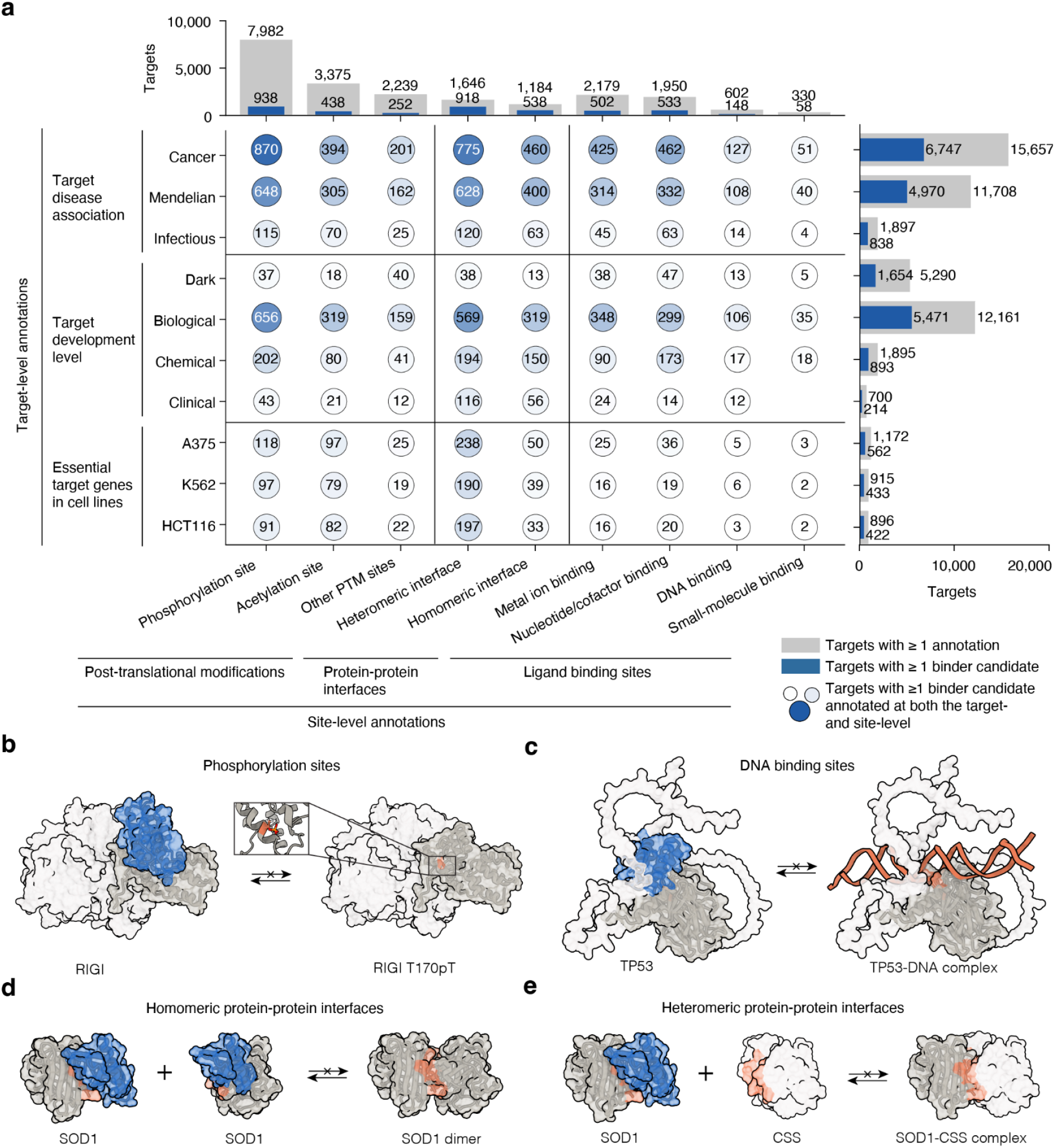
Functional landscape in proteins targeted with candidate binders. **a**, Landscape of functional perturbation hypotheses across target- and site-level annotations. Bubble matrix of intersections between target-level annotations (rows) and site-level annotations (columns). *Rows*: disease association (Cancer, Mendelian, Infectious; Pharos MONDO identifiers cross-referenced to MONDO disease categories), target development level (Pharos TDL: Dark, Biological, Chemical, Clinical) and gene essentiality (DepMap CRISPR knockout in A375, K562, HCT116). *Right bars*: per row, total annotated targets (grey) and the subset with ≥1 binder candidate (blue). *Columns*: PTM sites (UniProt), native protein-protein interfaces (PINDER) and ligand-binding sites (UniProt). *Top bars*: per column, annotated targets with at least one such site/interface annotation (grey) and the subset with ≥1 site/interface-perturbing binder candidate (blue). Bubble area and colour encode together with the overlaid count the number of targets that belong to the intersection of row and column annotation categories. Empty intersections are omitted. **b**, RIG-I (*RIGI*): a candidate whose epitope overlaps a regulatory phosphorylation site, nominating it as a perturbagen to block phosphorylation at that residue. **c**, TP53: a candidate whose epitope overlaps the DNA-binding surface, nominating it to occlude the protein-DNA interaction. **d**, **e**, SOD1: candidates whose epitopes overlap a homomeric (**d**, dimerization) or heteromeric (**e**, copper-chaperone CCS) protein-protein interface, nominating them to disrupt the respective interaction. PDB: SOD1 dimer, 3ECU; SOD1-CCS complex, 6FP6.

At the target level, binder candidates cover a substantial fraction of proteins annotated as cancer-, Mendelian disease-, or infectious disease-associated in Pharos and according to Mondo disease ontology. This shows that the Bindome may provide a number of affinity reagents that could be important to study disease-related processes. Coverage was also distributed across Pharos target development levels, including biological, chemical, clinical, and dark targets, suggesting that the Bindome provides candidate probes not only for well-studied proteins but also for less-characterized regions of the proteome. By intersecting UniProt target annotations with DepMap gene essentiality data, we found that approximately half of the genes classified as essential in each of three cancer cell lines, A375, K562, and HCT116, are targeted by at least one binder candidate.

At the epitope level, binder candidates frequently overlap with annotated functional sites. UniProt-annotated post-translational modification (PTM) sites were among the most common classes of potentially targetable sites, with phosphorylation and acetylation representing the largest categories. Using protein-protein interaction interfaces extracted from PINDER-curated structures, we found that approximately half of annotated homo- and heteromeric interfaces overlap at least one binder epitope for one of the interaction partners. This suggests that such reagents could be used to perturb specific protein interactions, which are very difficult to perturb selectively with chemical inhibitors or knockout studies. We also examined UniProt ligand-binding annotations and found that roughly one quarter of targets with annotated ligand-binding sites have at least one binder candidate whose epitope overlaps with those sites. Metal ion- and nucleotide/cofactor-binding sites were the most frequently annotated and most frequently overlapped categories, whereas DNA- and small-molecule-binding sites were less commonly targeted. Nevertheless, we provide binder candidates whose epitopes overlap annotated DNA-binding regions in 148 targets.

These annotations are not sufficient to ensure that the binders will perturb the function of the native protein. Instead they portray the breadth of experimentally testable hypotheses in which a binder may sterically occlude, stabilize, compete with, or otherwise modulate a functional site, interaction surface or target conformation. To illustrate how the Bindome can support such hypothesis generation, we examined intersections between target-level and epitope-level functional annotations, highlighting four representative examples (Fig. 3b-e).

For RIG-I/DDX58, a protein involved in antiviral RNA sensing, a binder candidate overlaps a region containing an annotated phosphorylation site. Such a candidate could be used to test whether engagement of this region alters phosphorylation-dependent regulation or downstream innate immune signalling (Fig. 3b). For TP53, a highly studied tumour-suppressor and transcription factor, a candidate overlaps the annotated DNA-binding region, suggesting a possible route to perturb DNA engagement or probe the structural accessibility of this functional surface (Fig. 3c). For SOD1, a Mendelian disease-associated protein implicated in amyotrophic lateral sclerosis (ALS), binder candidates overlap both homomeric and heteromeric protein-protein interfaces, including interfaces involved in SOD1 dimerization and interaction with its copper chaperone CCS (Fig. 3d,e). To facilitate the access of the Bindome to the wider scientific community we created a web resource described below.

### Interactive, programmatic and agentic access

All binder candidates are freely available through the Bindome website (https://bindome.epfl.ch). The resource provides three complementary modes of access: an interactive web interface for exploratory analysis, a standardized API for automated retrieval within analysis pipelines, and an MCP server for agentic querying through large language models (LLMs).

The web interface enables users to search the Bindome by target UniProt accession or entry name and retrieve all binder candidates designed against a given protein (Fig. 4a,e). Retrieved candidates can be ranked interactively using AF-derived confidence metrics and inspected in the context of the full-length target structure. A dedicated download section further enables retrieval of the complete database or predefined subsets of interest, such as binders predicted to disrupt protein-protein or DNA-protein interactions, occlude ligand- or metal-binding sites, or mask PTM sites. These downloads include the corresponding sequences, predicted structures, confidence metrics, and PAE matrices (Fig. 4b).

**Figure 4.**
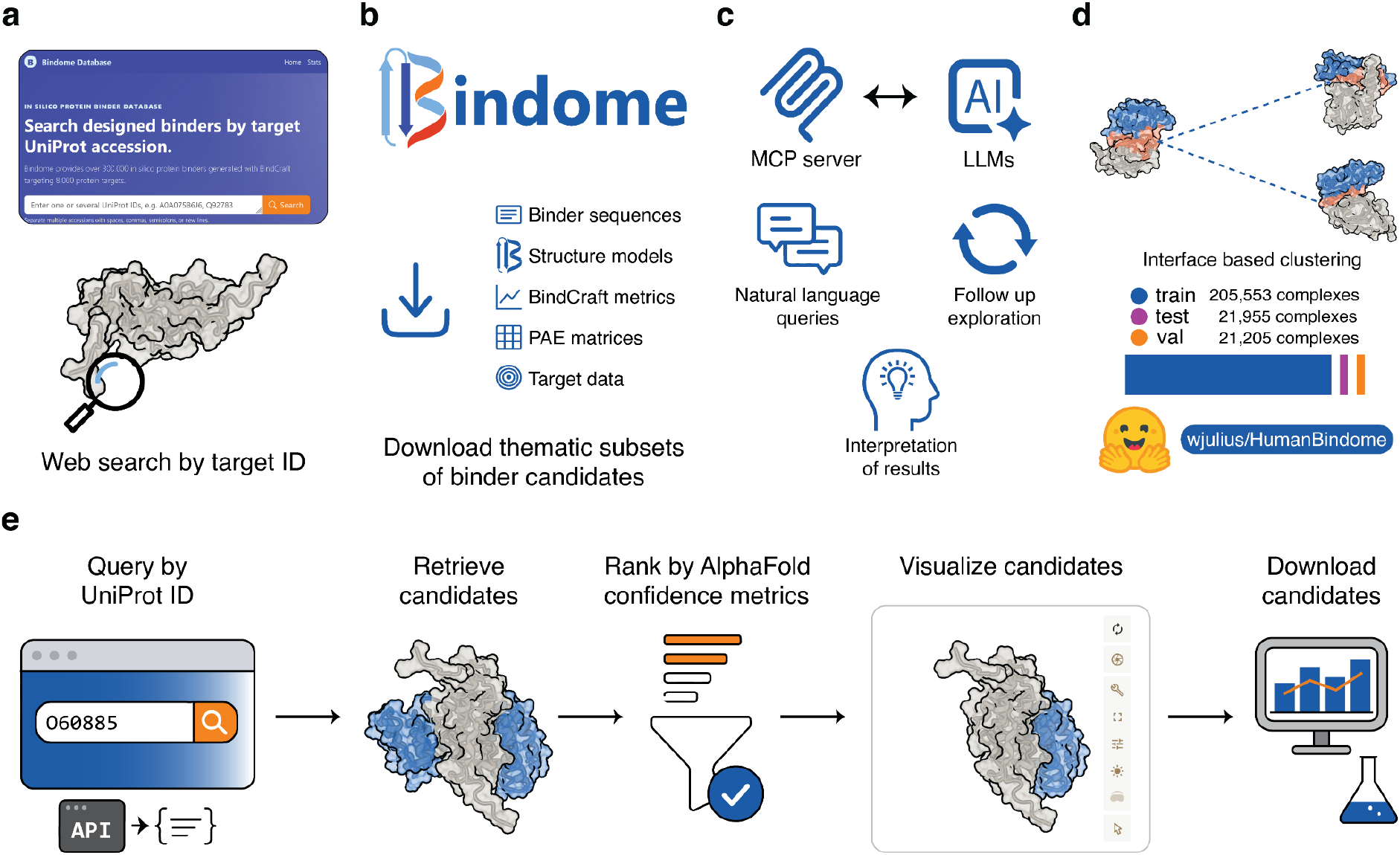
Bindome accessibility and programmatic interfaces. The Bindome’s *in silico* binder candidates are reachable through four complementary routes. **a**, The web interface (https://bindome.epfl.ch) enables target-based searches using UniProt accessions or entry names. **b**, Bulk download provides the complete dataset and large subsets with their corresponding target lists, BindCraft metrics, binder sequences, predicted structures and PAE matrices. **c**, An MCP server (https://bindome.epfl.ch/mcp) enables large language models to query the Bindome in natural language, interpret returned candidates and support iterative follow-up exploration. **d**, Leakage-controlled machine-learning splits, generated by interface-based clustering, are available as train, validation and test splits on Hugging Face (https://hf.co/datasets/wjulius/HumanBindome). **e**, Example web/API workflow: users query the Bindome with a target UniProt accession, retrieve the corresponding binder candidates, rank them using AlphaFold confidence metrics, inspect predicted complexes in the context of the full-length target structure and download selected results for downstream analysis.

For programmatic access, the Bindome exposes a 3D-Beacons API^37,38^, allowing binder candidates and associated metadata to be retrieved in a standardized and automated manner. This facilitates integration with external resources, including UniProt and PDBe-KB, and enables incorporation of the Bindome’s data into downstream computational workflows. Finally, the Bindome can also be queried through an MCP server, which connects the database directly to LLM-based interfaces (Fig. 4c). This allows users to ask questions in plain language rather than constructing explicit database queries. The model can then retrieve relevant candidates, summarize, and interpret the results, and support iterative refinement of the search within the same conversation. This agentic access mode broadens the usability of the Bindome, particularly for non-expert users seeking to explore or interpret binder candidates.

Given the demonstrated importance of distillation data for machine-learning model development^21,39^, we further made the Bindome dataset available on HuggingFace with leakage-controlled splits based on interface clustering. The splits comprise 205,553 training complexes, 21,205 validation complexes, and 21,955 test complexes (Fig. 4d).

## DISCUSSION

We created the human Bindome, a proteome-scale atlas of high-confidence *in silico* binder candidates. Its primary contribution is a new protein-based reagent resource, one that pairs broad proteome coverage with a defined sequence, a predicted binder-target structure with a defined epitope, and explicit confidence metrics for every binder candidate.

Existing affinity-reagent databases centre on antibodies and trade coverage against characterization. While antibody-based reagents and the databases that aggregate them have been crucial to biological research, their limitations have led to a situation severe enough to be described as an antibody reproducibility crisis^3^. The Bindome presents a novel type of designed candidate affinity reagent. Although these binder candidates still await experimental validation, they are extensively characterized *in silico*, with structural models that include the target side and freely available sequences for recombinant production. The Bindome’s overall proteome target coverage significantly exceeds that of databases offering comparable levels of *in silico* characterization.

The candidate binders are immediately suitable for detection applications such as labelling specific targets in microscopy or enrichment by affinity-based pull-down. The more far-reaching opportunity, however, is to use the candidate binders for perturbation in cellular contexts. Intracellular perturbation requires binders that are genetically encodable and stable in the reducing cytoplasm, which rules out full-length antibodies and their disulfide-dependent fragments in favour of small single-domain or de novo designed formats. Nanobodies proved such intrabodies can modulate targets in living cells but remain gated by per-target stability and one-at-a-time discovery^1,40^, a bottleneck that designed binders can in principle remove.

Genome- and transcriptome-wide screens have shown that systematic perturbations can reveal principles of cellular organization invisible to single-target experiments, but equivalent capabilities at the protein level have been limited by the lack of binders, which in turn was previously due to the low success rates and limited scalability of design and discovery methods. Because each of the Bindome’s candidates is a defined, genetically encodable sequence designed against a chosen target and a defined site, it provides a putative perturbagen that can be deployed to block interactions, modulate activity, alter localization, or recruit effectors^16^, and to do so by targeting specific epitopes rather than whole proteins. This opens systematic loss- and gain-of-function screens at proteome scale with site specific resolution, including against targets that lack tractable small-molecule pockets.

The underlying BindCraft pipeline has been validated in biochemical and functional assays, with binding success rates ranging from 10 to 100% depending on target^17^. Because the Bindome spans a far wider target diversity than has been tested, realized success rates across the atlas remain uncertain, and selectivity has yet to be characterized in depth. Selectivity is a twofold concern: cross-reactivity with unrelated proteins in the crowded cellular environment, and, more specifically, cross-reactivity among paralogs. The latter risk is heightened by our finding that binder candidates preferentially engage conserved epitopes (Fig. 2d) and by the family-level epitope convergence evident across CATH superfamilies (Fig. 2c). Both are preconditions for *in cellulo* perturbation.

How to evaluate *in silico* proteome-scale binder design is itself an open question. Target coverage, the fraction of proteins with at least one candidate, is the obvious metric and reaches 77.8% of what we provisionally define as the targetable proteome. However, from a perturbation standpoint, coverage of distinct epitopes may matter as much as coverage of targets; measured as a fraction of solvent-accessible surface area, coverage is substantially lower, indicating that additional progress in binder design methods is necessary. It also raises the question whether every interface is a priori tractable for binder design.

The Bindome is a living resource built to be findable, accessible, interoperable, and reusable with open source data, and freely accessible through interactive, programmatic, and agentic interfaces. As coverage extends across the human proteome, the same approach could be applied to generate proteome-wide binder resources for other model organisms commonly used in biological research.

We expect openly characterized *in silico* binders to increasingly complement experimentally derived reagents, furnishing labelling reagents and perturbagens for proteome-scale experiments with the site specificity to disentangle biological mechanisms and surface actionable hypotheses for modulating disease-relevant pathways.

## ONLINE METHODS

### Proteome-scale binder generation

#### Data

Reviewed *Homo sapiens* entries were retrieved from UniProtKB/Swiss-Prot^32^. For each accession, membrane-related annotations were extracted to determine membrane-embedded residue-ranges in the canonical sequence. Structural models and predicted aligned error (PAE) matrices were downloaded from AF Protein Structure Database (AFDB, v6)^23^. In case of isoforms, the isoform matching the canonical UniProt sequence was chosen.

#### Domain segmentation and annotation

Domain residue ranges were computed from PAE matrices using graph-based clustering (resolution = 1.0; PAE cutoff = 15 Å, implemented by the pae_to_domain^24^ tool by Tristan Croll). Range boundaries were extended by ±2 buffer residues, and gaps in the computed domain ranges were included when comprising ≤ 21 residues. Domain coordinates were extracted from full-length AFDB PDB files according to computed boundaries. For each domain, the following metrics were computed: length; mean pLDDT; DSSP secondary structure composition with BioPython (v1.86) DSSP module^41,42^; normalized radius of gyration^43^ (R_g,norm_), R_g,norm_ = R_g_ / N^1/3^, where R_g_ is the root-mean-square deviation of Cα positions from their geometric centroid and N is the number of residues, providing a size-independent measure of compactness; Cα contact density (contacts per residue within a radius of 8 Å). Membrane overlap was determined by intersecting domain residue ranges with UniProt-annotated membrane-embedded residues.

#### Target domain selection

Domains were retained if they satisfied all of the following: mean pLDDT ≥ 0.7; length 30-550 residues; R_g,norm_ ≤ 5.0; and Cα contact density ≥ 2.0. Domains annotated by DSSP as consisting exclusively of single helices, double helices, or loops were additionally excluded, as were entries with UniProt-AFDB sequence mismatches or any overlap with membrane-embedded residues.

#### Binder design

Binders were designed using an accelerated variant of BindCraft with default design and filtering settings established in the BindCraft publication^17^ and without hotspot guidance. To accelerate binder design to proteome scale, we (i) embedded BindCraft in a parallelization framework using independent SLURM array jobs, (ii) modified the code to avoid costly recompilation of ColabFold class instances of AF^44^ and ProteinMPNN^45,46^ by fixing the binder length to 90 residues, and (iii) leveraged a domain segmentation strategy that reduced target size and thereby accelerated the generation process. After the backpropagation and before the sequence redesign by ProteinMPNN, we superimposed the target domain and the binder onto the full target structure to ensure that binders were generated against accessible sites. Following superposition, we assessed steric clashes between the binder and the full target structure. Trajectories exhibiting clashes were rejected, except when clashing residues were of low confidence, defined as either a maximum pLDDT below 0.7 as reported in the model from the AF database or a symmetrized PAE greater than 15 Å as computed from the PAE matrices from the AF database. For each domain we generated up to 52 BindCraft accepted designs passing additionally the full target superposition criteria.

### Target coverage analysis

Targets, as defined by UniProt accessions, were partitioned by two nested criteria: solubility (soluble vs. membrane/insoluble, precomputed from the same UniProt annotations as for the target domain filtering) and, within each, targetability (present in a curated, filtered domain-level target set) and, within targetable targets, binder coverage (present in the designed-binder target set). Percentages reported for each category are relative to the combined soluble and insoluble proteome total.

### Molecular function classification of design targets

Each target protein was assigned to a broad molecular-function (MF) category derived from the Gene Ontology^25^ (GO). Human gene MF annotations (aspect F, excluding NOT-qualified entries) were obtained from the Functionome resource^47^ and mapped onto the generic GO-slim MF subset (go-basic.obo, format version: 1.2, data version: releases/2026-03-25) by breadth-first traversal of “is_a” and “part_of” relationships, assigning each GO term to its closest generic-slim ancestor; terms lacking a slim ancestor were labelled “other molecular function”. For each gene, the slim category with the largest annotation support was retained as its primary MF class. Genes that were not found in the databases were labelled as “not annotated”; those that were missing molecular function annotation were labelled as “unknown function”. To obtain a small set of biologically interpretable classes for display, related slim terms were manually consolidated into 13 categories: catalytic activity, binding activity, regulators, transcription factors, molecular transducer activity, transporters, structural proteins, molecular adaptors, molecular carriers, cytoskeletal motor activity, and sequestering/preventing-interaction activity, plus not-annotated, and unknown function. Genes initially labelled “other molecular function” were further resolved by keyword matching (“binding,” “factor”) against their constituent GO term names and reassigned to the binding-activity or transcription-factor class accordingly. Those that could not be resolved were kept as “other molecular function”. This procedure produced a proteome-wide UniProt-to-function-category map.

### Comparison to affinity reagent databases

To contextualise the Bindome’s target coverage, we assessed four publicly available affinity reagent databases. For direct comparability with Bindome, we restricted each database to Swiss-Prot annotated human targets.

#### Antibodypedia

The coverage of Antibodypedia^6^ (https://www.antibodypedia.com; gene list accessed 15.06.2026) was assessed via direct gene-name matching of their gene list against Swiss-Prot primary gene symbols, and retrieval of all Swiss-Prot entries carrying an Antibodypedia cross-reference (“database:antibodypedia”). To avoid case-sensitivity artefacts, all gene symbols were uppercased prior to matching.

#### ABCD

Human gene targets from ABCD^7^ (AntiBodies Chemically Defined, https://web.expasy.org/abcd/; Version 17.0 (March 2026); accessed 03.06.2026) were matched to Swiss-Prot primary gene symbols (uppercase) to obtain the coverage.

#### Biocompare

Antigen names from Biocompare^48^ (https://www.biocompare.com/Antibodies/; accessed 03.06.2026) were resolved to Swiss-Prot accessions via the API (queried with scopes=symbol,alias,name, species 9606, score threshold ≥ 5.0).

#### SAbDab

To retain only entries in which both “antigen_species” and “heavy_species” contained “homo sapiens”, SAbDab^5^ (Structural Antibody Database, https://sabdab.opig.stats.ox.ac.uk/; accessed 12.06.2026) was filtered. PDB identifiers from these human-on-human structures were mapped to UniProt accessions via the SIFTS flat file^49,50^ (pdb_chain_uniprot.tsv.gz, EBI FTP, accessed 12.06.2026).

### Proteome surface coverage analysis

Per residue solvent accessible surface area^51^ (SASA) was calculated for each protein in the human proteome from its full length AFDB structure using the freesasa (v.2.0.3) Python package^52^. SASA was computed per atom and summed over each residue’s constituent atoms to obtain per residue values.

To evaluate binder induced coverage in this full length context, the binder-domain complex was superimposed onto the full length target by alignment of the shared Cα atoms of the cropped domain. This aligns the binder along with the domain in the full length structure.

Binder buried area was then computed per residue as the difference between the SASA of the unbound full length target and the SASA of that full length target with the binder superimposed. Consequently, the total accessible proteome surface includes only residues that are exposed in the full-length structure. Surfaces occluded by other regions of the same polypeptide chain, including interdomain interfaces, are therefore excluded, even if they are exposed in the isolated cropped domain. Where multiple binders targeted the same protein, each residue was assigned its maximum burial across binders and counted as covered if buried by at least one of them. Proteome surface coverage was then defined as the total binder covered surface area divided by the total accessible proteome surface.

### CATH-domain coverage analysis

#### Annotation

CATH superfamily annotations were assigned by structurally searching each human target domain against a local CATH Foldseek database^26,28^ using Foldseek easy-search. Searches were performed with high sensitivity (-s 9.5), an E-value threshold of 1e-3, up to 1000 prefilter hits per target domain, TM-align-based final alignment (--alignment-type 1), and no search-level coverage threshold (-c 0.0, --cov-mode 2). Hits were filtered to retain matches with query-normalized TM-score ≥ 0.5, where the query corresponds to the human target domain. Each human target domain with at least one passing CATH match was assigned to the best remaining CATH reference domain, ranked by the minimum of the query and reference CATH domain normalized TM-scores.

#### Surface projection of binder-targeted sites across four CATH superfamilies

Human proteins containing CATH superfamilies 3.30.200.20 for kinases, 2.130.10.10 for WD40 domains, and 1.20.920.10 for bromodomains were identified based on the annotation described above. For visualization, one representative target structure was selected for each family: CDK1 (P06493) for kinases, WDR5 (P61964) for WD40 domains, and BRD4 (O60885) for bromodomains. For each target-binder complex, the target chain was structurally aligned to the corresponding representative target structure by matching Cα atoms based on local sequence similarity, followed by iterative trimmed-core structural superposition. Complexes were retained only if the trimmed core alignment contained at least 25 Cα pairs and satisfied a maximum Cα-RMSD of 6.0 Å. For retained complexes, binder-targeted residues were identified as target residues making heavy-atom contacts within 4.0 Å of the binder. Contacted residues were then projected onto the representative structure by assigning them to the nearest representative residue after target-chain alignment, using a maximum projection distance of 7.0 Å. Counts were normalized by the maximum residue count within each representative structure.

### Epitope evolutionary conservation analysis

#### Multiple sequence alignments and per-residue evolutionary rates

For each target, an MSA was retrieved from the AFDB (v6) MSA endpoint and converted to a flat FASTA alignment with HH-suite^53^ (v3.3.0) reformat.pl, so that alignment column i corresponds to residue i of the full-length UniProt sequence. Per-domain sub-alignments were extracted at the domain’s AFDB residue positions. Evolutionary rates were computed per domain with Rate4Site^54,55^ (GitHub: https://github.com/pegerto/pyrate4site, v0.1.0) using the query as reference. Domains with a reference ≥3,000 residues or fewer than 2 usable sequences were excluded. MSAs deeper than 200 sequences were downsampled deterministically: exact duplicates were collapsed, the reference retained, and the remainder binned into 20 fractional-identity bins (over non-gap positions) and drawn round-robin from low to high identity to preserve diversity.

#### Epitope definitions and evolutionary-rate controls

In each target-binder complex, surface residues were defined by relative solvent accessibility (RSA = SASA/MaxASA) > 0.25, where SASA was computed per residue using Biopython’s Shrake-Rupley algorithm^56^ (v1.86) and maximum-ASA values from Tien et al.^41,57^. A binder’s epitope was defined as the surface residues with any atom within 4.0 Å of a binder atom. Epitopes were pooled across all of a domain’s binders to define consensus epitopes: residues contacted by any binder, by ≥30%, or by ≥90% of all the binders targeting each domain. For each consensus epitope, the mean Rate4Site rate was compared to a residue-number-matched control drawn without replacement from the domain’s surface-residue pool (a matched-background control that may include consensus-epitope residues).

### Bindome comparison to native protein-protein interactions

#### Data

Designed binder-target complexes were drawn from the Bindome, and native complexes from PINDER^29^ (release 2024-02, accessed via the pinder python package v0.5.0). For the Bindome, all binder-target complex structures were deduplicated at the level of target domain: for each target domain, the single binder with the highest average binder pLDDT was retained, with average interaction pTM used as a tiebreaker, yielding one representative design per target domain. For PINDER, dimeric assemblies were first filtered to biologically relevant, high-confidence interfaces (resolution ≤ 3.5 Å; interface label “BIO”; predicted interface probability ≥ 0.5; no interface-proximal atom gaps or missing interface residues within 4 Å), matched to a structure file by exact PINDER identifier, and then deduplicated at the level of the interacting UniProt pair: for each unique UniProt-UniProt interaction, the entry with the highest probability was kept, with lower crystallographic resolution used as a tiebreaker.

#### Interface size

For every retained complex, the SASA was computed with the Shrake-Rupley algorithm in Biopython (v1.86) at the atomic level, separately for the complex and for each partner chain in isolation. For each atom in an isolated chain, the buried SASA was defined as buried(atom) = max(0, SASA_isolated_ − SASA_complex_), and residue-level buried SASA was obtained by summing buried SASA over all atoms of a residue. A residue was classified as an interface residue if its buried SASA exceeded a small numerical threshold (10⁻⁶ Å², to exclude floating-point noise). The interface area was reported as half the total residue-level buried SASA summed over both partners.

#### Interface polarity

Interface residues (as defined above) were further classified as polar if their residue type belonged to the union of polar-uncharged (Ser, Thr, Asn, Gln, Tyr, Cys), acidic (Asp, Glu), and basic (Lys, Arg, His) amino acids. For each complex, the polar interface fraction of a given chain was computed as the number of polar interface residues divided by the total number of interface residues on that chain. For the Bindome data analysis, the binder-side polar fraction was used, thereby reflecting the interface composition of the designed binder. For PINDER, polar fractions were computed for both partner chains and pooled, so that the native distribution reflects polar character on either side of a native interface.

#### Secondary structure interface combinations

Per-residue secondary structure was assigned with BioPython (v1.86) DSSP module on each complex structure, and the eight-state DSSP code was collapsed into three classes: helix (H, G, I), sheet (E, B), and loop (all remaining codes). For each chain in each complex, interface residues were binned into these three classes, and the predominant secondary-structure class of that chain’s interface was defined as the class with the highest interface-residue count.

#### Fold novelty

To assess the structural novelty of designed binder folds relative to known and predicted protein structures, we performed a structural similarity search using Foldseek easy-search (commit 718d421) against the AFDB. Binder monomer prediction models were used as the query database, and a local Foldseek-indexed copy of the AFDB (v6 2025_03) served as the target database. Searches were run using the combined 3Di/amino-acid local alignment mode (--alignment-type 2), with coverage computed relative to the query length (--cov-mode 2) and no minimum coverage threshold applied (-c 0.0). All other Foldseek parameters were left at default values. For each binder, the highest-scoring hit according to the query TM-score against the AFDB was used to evaluate whether the designed fold corresponded to a previously observed structural topology or represented a novel fold.

#### Interface novelty

To assess the structural novelty of the designed binder-target interfaces relative to experimentally determined protein-protein interfaces, we performed an interface similarity search against PPIRef using the PPIRef toolkit^30^. For each designed complex, the interface between the natural target and the designed binder was extracted using a 6 Å heavy-atom contact cutoff, matching the interface definition used to construct the PPIRef reference set of dimeric interfaces from the PDB. Because exhaustively aligning each interface against all of PPIRef is computationally prohibitive, candidate matches were first retrieved with iDist^30^, retaining reference interfaces within an iDist distance of 0.1 of each query. The retrieved up to 10 candidates were then superposed onto the query with US-align^58^ (version 20260527) in multimeric alignment mode (-mm 1 -ter 1), and the interface TM-score was taken as the higher of the two length-normalised scores. For each binder, novelty was summarised by its best interface TM-score across all retrieved candidates. All other iDist and US-align parameters were left at default values.

#### Induced-fit signal from structural comparison

To quantify the conformational change induced upon binding, we compared predicted unbound and bound complex models for each binder-target pair and computed the Cα interface RMSD after superposition. Two RMSD distributions were assessed: (i) the binder signal, defined as the Cα RMSD between the binder’s unbound and bound conformations at the binding interface, and (ii) the target signal, defined analogously for the target protein.

### Coverage of allosteric sites

Human allosteric target and modulator annotations were taken from the Allosteric Database^31^ (ASD, release 2023-09). For each ASD record on a Bindome-covered target, site-lining residues were derived from the deposited structure. Target contact residues were defined by a heavy-atom distance cutoff specific to modulator type: 4.0 Å for protein, peptide, or ion modulators and 5.0 Å for small-molecule or glycan ligands. Contact residues were converted from author numbering to canonical UniProt numbering via the mmCIF SIFTS annotation. Designed-binder interfaces were defined analogously, using the same 4.0 Å heavy-atom cutoff. A covered target was scored as having a confirmed overlap if the ASD contact residues shared at least one residue with the designed-binder interface residues.

### Proteome-scale nomination of perturbagen candidates

#### Bindome

For every target-binder model, interface residues were identified as standard-residue heavy atoms in cross-chain contact within 4.0 Å.

#### Target development level and disease association

Target development level (Tclin, Tchem, Tbio, Tdark; shown in the figure as Clinical, Chemical, Biological, and Dark, respectively) and associated MONDO^34^ disease identifiers were retrieved for every target from the Pharos^33^ GraphQL API (access 28 May 2026). Disease-category membership (cancer, infectious, Mendelian/hereditary and rare) was assigned based on the MONDO disease ontology (release 2026-05-05, accessed 28 May 2026). Each target was assigned to a category if its Pharos-linked disease annotations fell under that category in the ontology.

#### Target essentiality

Cell-line-specific essential genes were derived from DepMap^36^ Public 26Q1 CRISPR gene-effect Chronos scores for HCT116, K562 and A375, calling a gene essential in a given line when its gene-effect score was below −1.0. Gene symbols were mapped to UniProt accessions through the reviewed human proteome; ambiguous symbol-to-accession mappings were resolved by a first-listed-alias priority rule and otherwise discarded.

#### Allosteric-site coverage

Targets with at least one annotated allosteric site were fetched from the human entries of the Allosteric Database^31^ (2023-09 release). Because this release encodes allosteric- and ligand-site positions as PDB and ligand identifiers rather than canonical protein residue numbers, allosteric sites were not resolved to residues or intersected with binder interfaces; allosteric-site coverage is therefore reported only as a whole-target (UniProt accession) annotation.

#### Post-translational modification and ligand binding site annotations

Perturbation-relevant site and region annotations were retrieved for every target accession from the UniProtKB^32^ REST API (access: May 30, 2026). To avoid misclassification, raw feature descriptions were mapped to a controlled vocabulary using exact-match allowlists. For each annotated feature we tested whether a binder’s epitope residue set overlapped with the annotated residue or by at least one residue with the annotated residue range. For the bubble matrix, the PTM classes were aggregated into phosphorylation, acetylation, and a pooled “other” category. The “other” category is an explicit allowlist of modification types (including methylation, succinylation, ADP-ribosylation, lipid modifications such as palmitoylation and myristoylation, sulfation, hydroxylation, and related redox modifications) and is therefore not exhaustive. A number of rarer modification types present in the UniProt annotations were excluded from analysis due to their low frequency. Glycosylation was excluded due to limited perturbation potential. Ligand-binding features were grouped into the four classes above (nucleotide/cofactor, metal ion, DNA, and other small molecules).

#### Protein-protein interfaces

Experimentally observed human-human interfaces were taken from PINDER^29^ (2024-02 release, accessed via the pinder python package v0.5.0), restricted to high-confidence biological assemblies (resolution ≤ 3.5 Å; PRODIGY-cryst biological label with probability ≥ 0.5; no interface residues or atoms missing within 4 Å). Receptor- and ligand-side interface residues were computed at the same 4.0 Å radius and mapped to UniProt positions, and each interface was classified as homomeric or heteromeric from the PINDER complex-type annotation. A binder was counted as disrupting an interface when its mapped positions overlapped the corresponding PINDER interface residues for that accession by at least one residue.

#### Bindome Server and API

The Bindome’s data are hosted at EPFL and exposed through a 3D-Beacons API^37,38^. The database was integrated into the 3D-Beacons framework to provide standardized programmatic access and enable interoperability with UniProt-linked resources and downstream tools.

### Leakage-controlled data splits for machine learning application

#### Overview

Leakage-controlled train, validation, and test splits were generated by adapting the PINDER splitting strategy for protein-protein interaction benchmarks^29^ to the Bindome target-binder dataset. Each of the Bindome’s entries was treated as a bound two-chain complex composed of a target chain and a binder chain. Components of the original PINDER workflow that are specific to experimental structural datasets, including prioritization by experimental method, crystallographic resolution and biological-assembly confidence, were not applied.

#### Clustering

Interface residues were defined separately on the two chains using the same backbone-contact convention as PINDER. Complete target and binder chains were aligned all-vs-all with Foldseek. An edge between two chains was retained only when the Foldseek-aligned region covered ≥50% of an interface residue set, so that graph construction reflected similarity of the binding interface rather than global structural similarity alone. The resulting interface-similarity graph was thresholded with the PINDER default alignment-lDDT threshold of 0.70. Weighted asynchronous label propagation was used to assign chain-interface cluster labels. Each designed dimer was then assigned to a paired-interface cluster ({C_target, C_binder}) from the two participating chain-interface cluster identifiers.

#### Split and deduplication

Train, validation, and test assignment was performed at the paired-interface cluster level, such that complexes within the same paired-interface cluster were assigned to the same partition. After holdout clusters (validation and test) were selected, we applied PINDER’s depth-2 transitive deleaking procedure to remove additional training-side complexes connected to holdout systems through the Foldseek/interface graph. In this process, candidate training complexes connected to holdout complexes through one or two edges in either the Foldseek interface-similarity graph (using the PINDER default deleaking edge threshold of 0.55) or a separately constructed interface-filtered MMseqs sequence-similarity graph, retaining only whole-chain sequence alignments spanning at least 50% of an interface residue set, were assigned to the deleaked partition and excluded from the usable benchmark.

#### Summary and audit

This procedure produced a leakage-controlled benchmark of 248,713 target-binder complexes, corresponding to 81.2% of the input dataset. The final partitions contain 205,553 training complexes, 21,205 validation complexes and 21,955 test complexes, spanning 127,743, 10,140, and 10,139 paired-interface clusters, respectively. As an independent redundancy audit, we examined ProteinMPNN sequence variants generated from the same binder backbone, which were not provided to the clustering procedure. Among 120,609 backbones with sister variants, 120,082 remained within a single paired-interface cluster, indicating that 99.56% of these redundant design groups were captured by the interface-based clustering. After deleaking, only 23 of the 88,720 multi-member backbone groups in the usable benchmark (0.026%) had sister variants split between the training and held-out partitions.

#### Use of large language models

Large language models were used to assist with code development, drafting of the manuscript and brainstorming. All AI-assisted outputs were reviewed and verified by the authors, who take responsibility for the content, code and its accuracy.

## Data Availability

All of the Bindome’s entries are openly accessible through a web interface (https://bindome.epfl.ch), an API (https://bindome.epfl.ch/docs), and an MCP server (https://bindome.epfl.ch/mcp), and binders generated in the future will be added to this database. Binder data and machine-learning splits are additionally available for download as a Hugging Face dataset (https://hf.co/datasets/wjulius/HumanBindome). All binder data are released under a Creative Commons Attribution 4.0 International (CC BY 4.0) license, permitting both academic and commercial use with attribution.

## Code Availability

All code for the generation of the Bindome’s binder candidates is available in a GitHub repository (https://github.com/wejulius/Bindome).

## Acknowledgements

We acknowledge the EuroHPC for awarding this project access to the EuroHPC supercomputer Leonardo hosted by CINECA in Italy (EHPC-AIF-2025SC03-099). J.W. was supported by a SwissAI PhD Fellowship 2025. We also thank EMBO for the Scientific Exchange Grant (n° 12335) to support R.H.’s stay at EPFL. We thank Flávia Mayumi Odahara de Abreu for designing the Bindome logo. Further, we thank the members of the Correia, Ablasser, Taipale and Winter labs for discussions.

## Author Contributions

J.W., A.D., and B.C. conceived the study and acquired funding. J.W. developed the pipeline and performed the binder generation, with contributions from A.D., S.G., K.T., M.T., M.H., G.W., and B.C.. A.D. built the web server, API, and MCP, with contributions from R.P., D.D., J.W., D.S., S.N., M.A., J.F., S.V., and B.C. The results were analysed by J.W., A.D., J.K., A.B., R.H., E.E., and B.C.. J.W., A.D., J.K., A.B., and B.C. wrote the manuscript with input from all authors. All authors reviewed and approved the final manuscript.

## Extended Data

**Extended Data Fig. 1.**
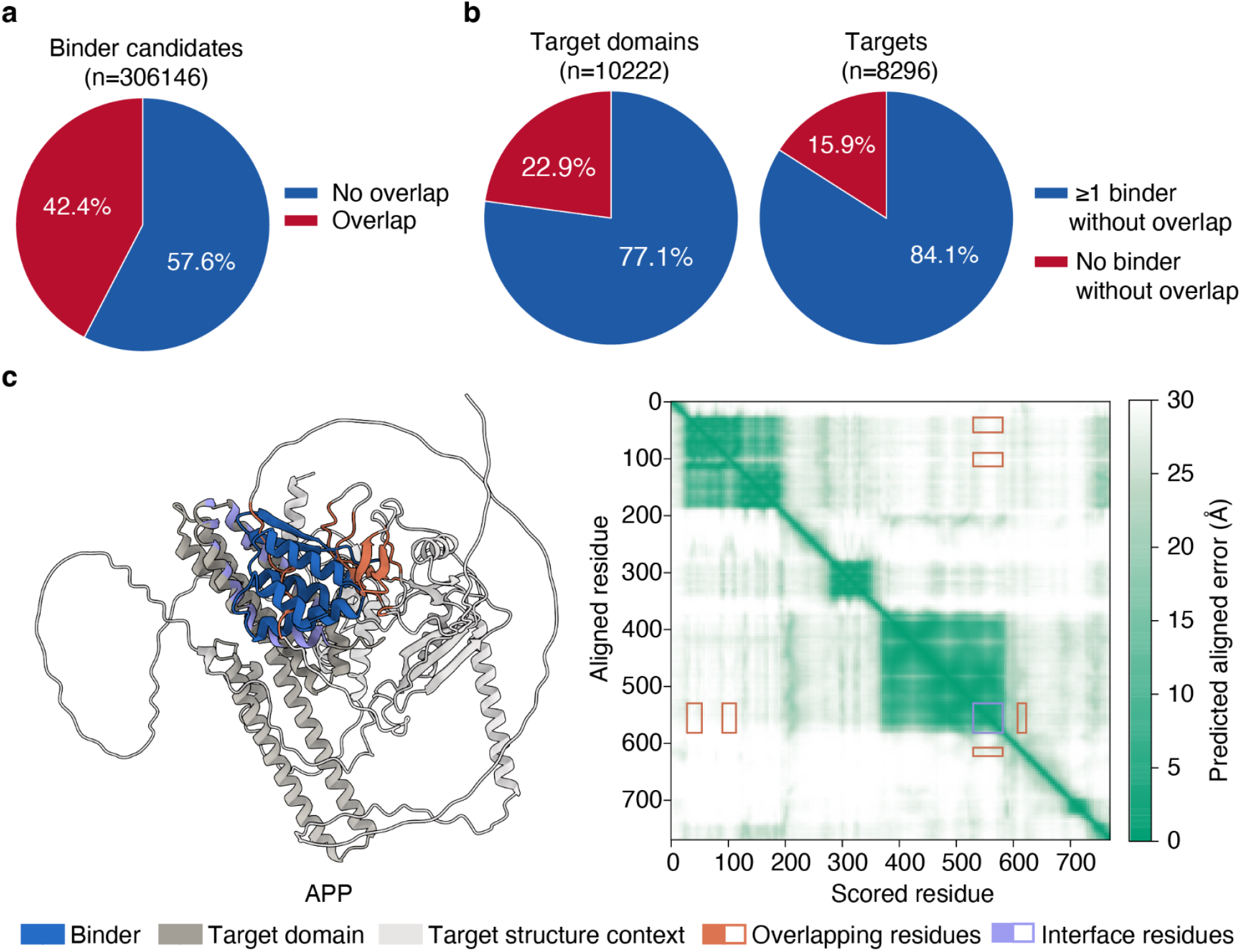
Overlap between designed binders and the full-length target. **a,** Binder candidates classified by whether they overlap the target after superimposing their target domain onto the full-length target structure. **b,** Target domains and full-length targets, categorized by whether all their binders overlap or at least one binder is non-overlapping. **c,** Representative target structure with a binder that overlaps the full-length target yet was retained, owing to low predicted aligned error (PAE) over the overlapping target region.

**Extended Data Fig. 2.**
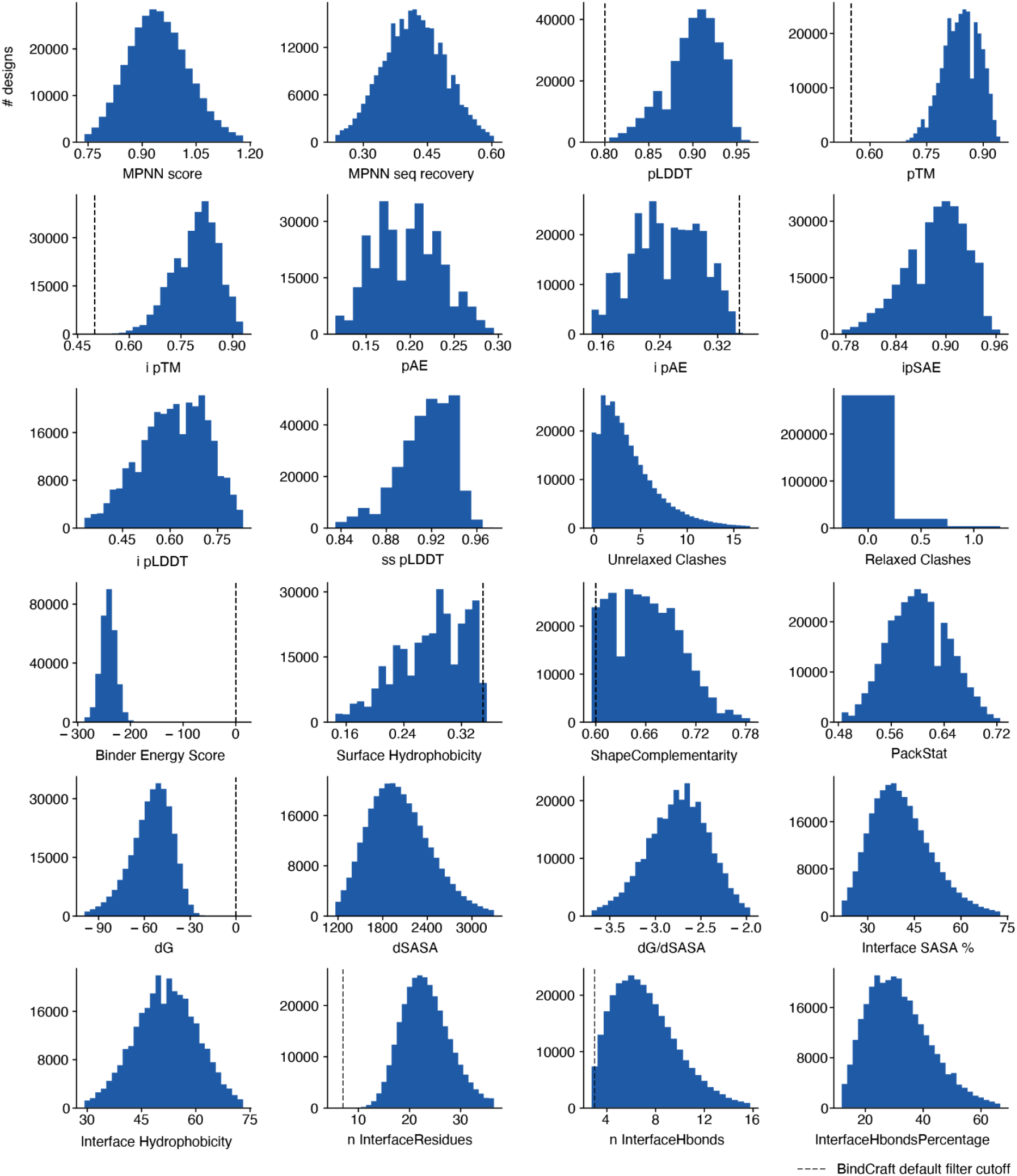
Distributions of BindCraft design metrics across all of the Bindome’s designs (1 of 2). Histograms show the per-design metric distributions. Dashed lines mark BindCraft default filter cutoffs where applicable. Panels are split across Extended Data Figs. 1 and 2 for layout only.

**Extended Data Fig. 3.**
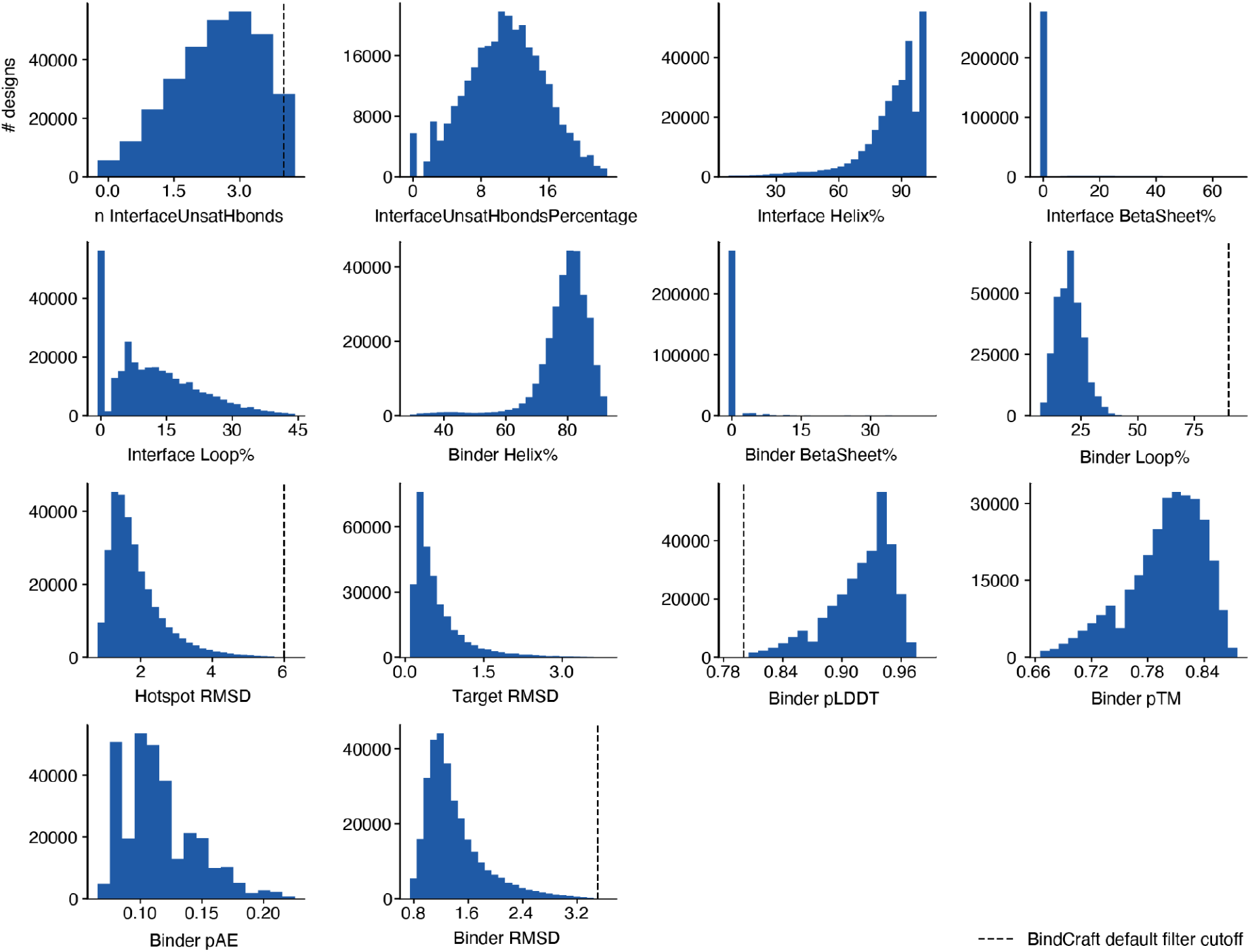
Distributions of BindCraft design metrics across all of the Bindome’s designs (2 of 2). Continuation of Extended Data Fig. 1. See that figure for full legend.

**Extended Data Fig. 4.**
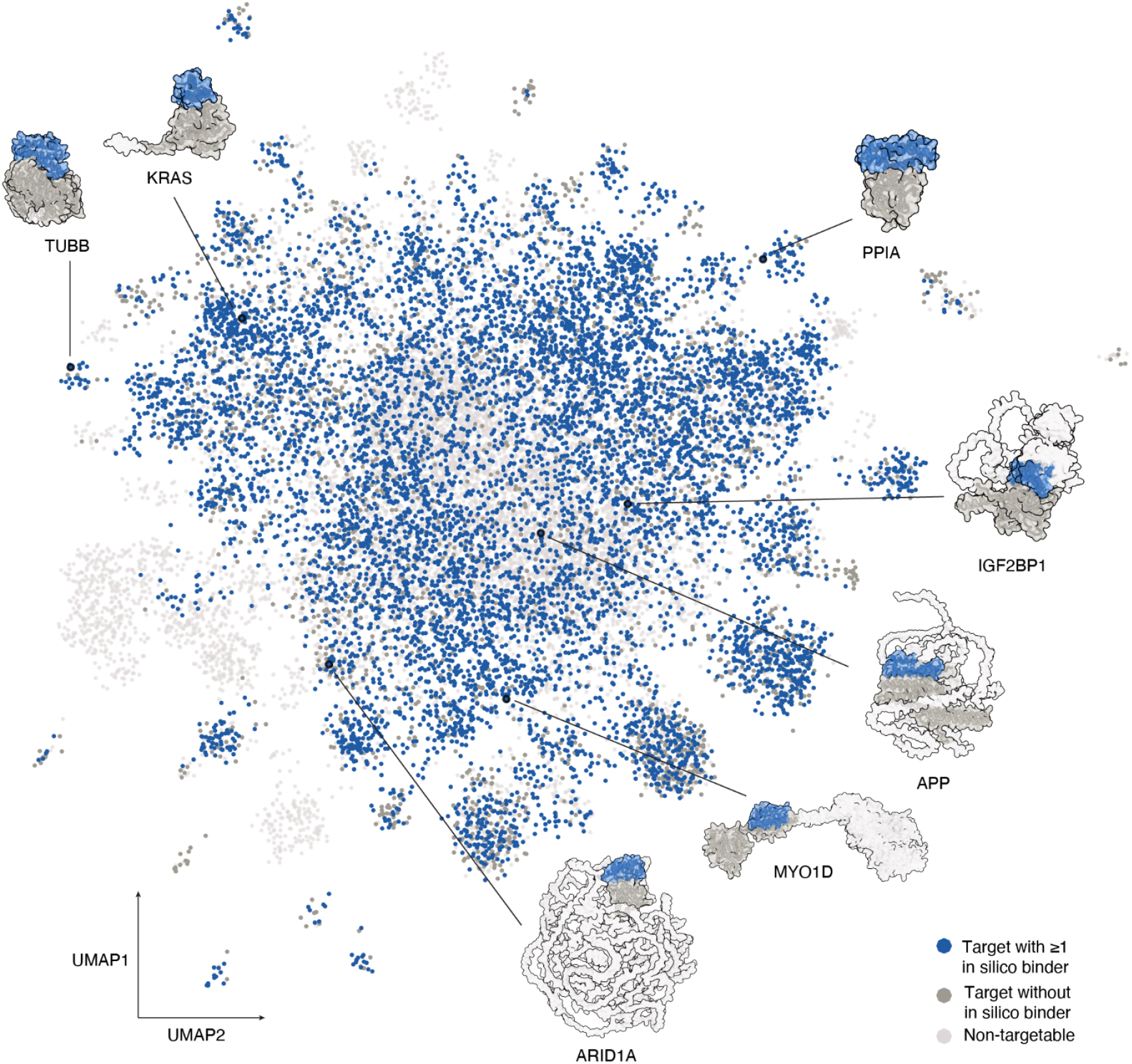
Structural UMAP of the human proteome with targets that are addressed in the Bindome highlighted. UMAP projection of human proteome structures derived from all-versus-all TM-score comparisons, with representative target-binder complexes annotated: KRAS, a compact oncogenic GTPase; TUBB, a globular cytoskeletal protein; MYO1D, a multidomain motor protein; APP, a membrane-associated Alzheimer’s disease protein; ARID1A, a large chromatin-remodelling tumour suppressor with extensive predicted disorder; and IGF2BP1, a multidomain RNA-binding protein.

**Extended Data Fig. 5.**
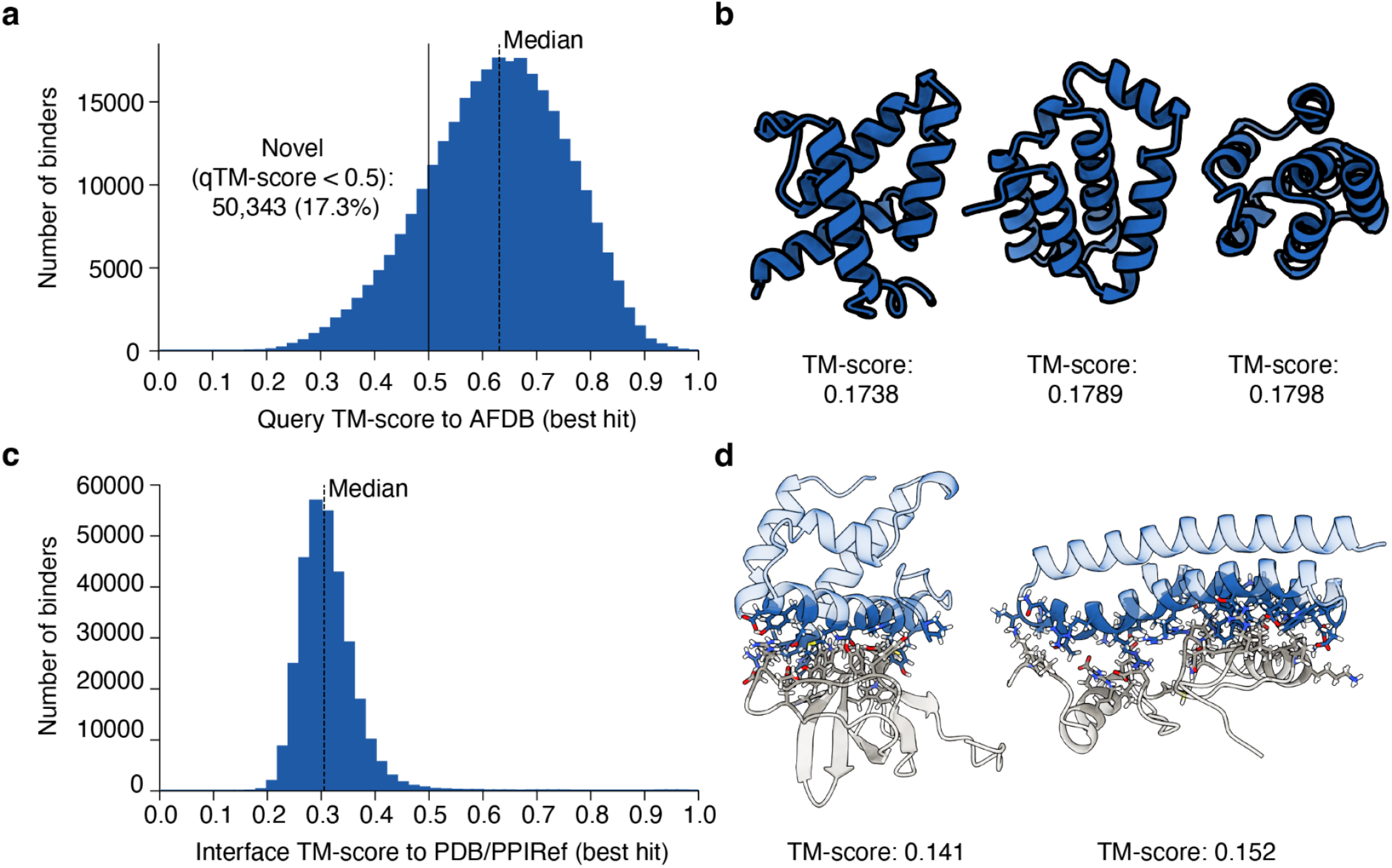
Structural and interface novelty of designed binders. **a,** Fold novelty analysis. A Foldseek search against the AlphaFold Database (AFDB) shows that 17.3% of monomer binders adopt a fold with a maximum qTM-score < 0.5 to any AFDB structure, indicating a novel fold. **b,** Representative binders with such novel folds. **c,** Interface novelty analysis. **d,** Representative binders with novel interfaces.

**Extended Data Fig. 6.**
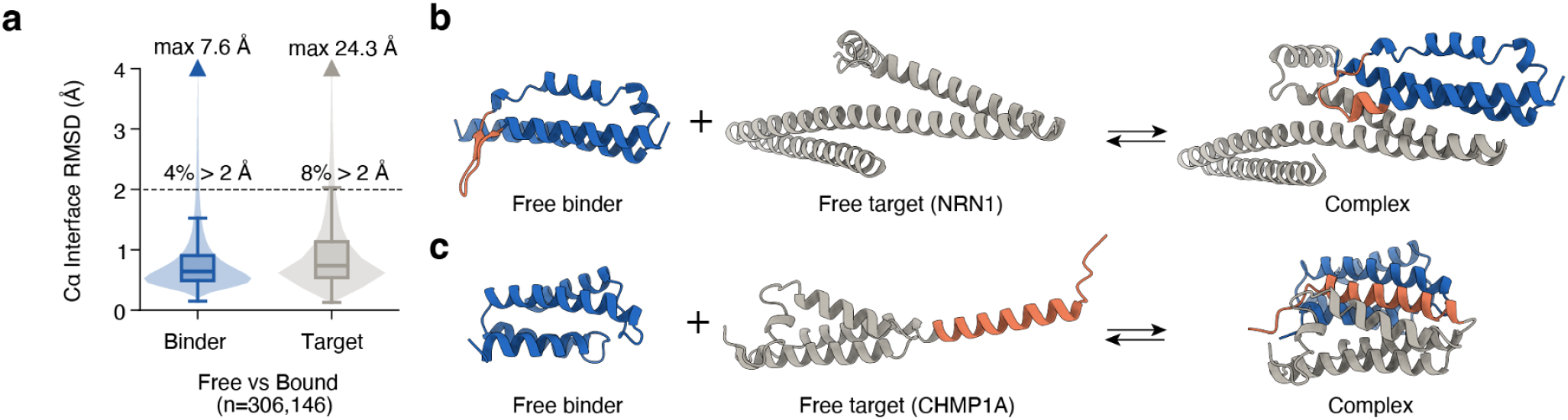
Induced fit. **a**, Prevalence of induced fit on the binder and target sides. **b**, Representative example of induced fit on the binder side. **c**, Representative example of induced fit on the target side.

## References

1. Helma, J., Cardoso, M. C., Muyldermans, S. & Leonhardt, H. Nanobodies and recombinant binders in cell biology. J. Cell Biol. 209, 633–644 (2015).

2. Garisto, D. Science sleuths uncover more than 100 suspicious images in Thermo Fisher antibody catalogue. Nature 654, 303–304 (2026).

3. Baker, M. Reproducibility crisis: Blame it on the antibodies. Nature 521, 274–276 (2015).

4. Ayoubi, R. et al. Scaling of an antibody validation procedure enables quantification of antibody performance in major research applications. eLife 12, RP91645 (2023).

5. Dunbar, J. et al. SAbDab: the structural antibody database. Nucleic Acids Res. 42, D1140–D1146 (2014).

6. Björling, E. & Uhlén, M. Antibodypedia, a Portal for Sharing Antibody and Antigen Validation Data. Mol. Cell. Proteomics 7, 2028–2037 (2008).

7. Lima, W. C. et al. The ABCD database: a repository for chemically defined antibodies. Nucleic Acids Res. 48, D261–D264 (2020).

8. Freedman, L. P., Cockburn, I. M. & Simcoe, T. S. The Economics of Reproducibility in Preclinical Research. PLOS Biol. 13, e1002165 (2015).

9. St Johnston, D. The art and design of genetic screens: Drosophila melanogaster. Nat. Rev. Genet. 3, 176–188 (2002).

10. Boone, C., Bussey, H. & Andrews, B. J. Exploring genetic interactions and networks with yeast. Nat. Rev. Genet. 8, 437–449 (2007).

11. Shalem, O., Sanjana, N. E. & Zhang, F. High-throughput functional genomics using CRISPR–Cas9. Nat. Rev. Genet. 16, 299–311 (2015).

12. Platt, R. J. et al. CRISPR-Cas9 Knockin Mice for Genome Editing and Cancer Modeling. Cell 159, 440–455 (2014).

13. Roth, T. L. et al. Pooled Knockin Targeting for Genome Engineering of Cellular Immunotherapies. Cell 181, 728–744.e21 (2020).

14. Lue, N. Z. & Liau, B. B. Base editor screens for *in situ* mutational scanning at scale. Mol. Cell 83, 2167–2187 (2023).

15. Kondo, S. & Perrimon, N. A genome-wide RNAi screen identifies core components of the G₂-M DNA damage checkpoint. Sci. Signal. 4, rs1 (2011).

16. Suiter, C. C. et al. Multiplex design and discovery of proximity handles for programmable proteome editing. 2025.10.13.681693 Preprint at 10.1101/2025.10.13.681693 (2025).

17. Pacesa, M. et al. BindCraft: one-shot design of functional protein binders. 2024.09.30.615802 Preprint at 10.1101/2024.09.30.615802 (2025).

18. Stark, H. et al. BoltzGen: Toward Universal Binder Design. 2025.11.20.689494 Preprint at 10.1101/2025.11.20.689494 (2025).

19. Watson, J. L. et al. De novo design of protein structure and function with RFdiffusion. Nature 620, 1089–1100 (2023).

20. Butcher, J. et al. De novo Design of All-atom Biomolecular Interactions with RFdiffusion3. 2025.09.18.676967 Preprint at 10.1101/2025.09.18.676967 (2025).

21. Didi, K., et al. Scaling Atomistic Protein Binder Design with Generative Pretraining and Test-Time Compute. Preprint at 10.48550/arXiv.2603.27950 (2026).

22. Balbi, P. E. M., et al. Mapping targetable sites on the human surfaceome for the design of novel binders. Proc. Natl. Acad. Sci. 123, e2506269123 (2026).

23. Varadi, M. et al. AlphaFold Protein Structure Database: massively expanding the structural coverage of protein-sequence space with high-accuracy models. Nucleic Acids Res. 50, D439–D444 (2022).

24. Croll, T. tristanic/pae_to_domains. https://github.com/tristanic/pae_to_domains (2026).

25. The Gene Ontology knowledgebase in 2026. Nucleic Acids Res. 54, D1779–D1792 (2025).

26. Knudsen, M. & Wiuf, C. The CATH database. Hum. Genomics 4, 207 (2010).

27. Waman, V. P. et al. CATH v4.4: major expansion of CATH by experimental and predicted structural data. Nucleic Acids Res. 53, D348–D355 (2025).

28. van Kempen, M. et al. Fast and accurate protein structure search with Foldseek. Nat. Biotechnol. 42, 243–246 (2024).

29. Kovtun, D. et al. PINDER: The protein interaction dataset and evaluation resource. 2024.07.17.603980 Preprint at 10.1101/2024.07.17.603980 (2024).

30. Bushuiev, A., et al. Learning to design protein-protein interactions with enhanced generalization. Preprint at 10.48550/arXiv.2310.18515 (2024).

31. Huang, Z. et al. ASD: a comprehensive database of allosteric proteins and modulators. Nucleic Acids Res. 39, D663–D669 (2011).

32. The UniProt Consortium. UniProt: the Universal Protein Knowledgebase in 2023. Nucleic Acids Res. 51, D523–D531 (2023).

33. Kelleher, K. J. et al. Pharos 2023: an integrated resource for the understudied human proteome. Nucleic Acids Res. 51, D1405–D1416 (2023).

34. Vasilevsky, N. A. et al. Mondo: integrating disease terminology across communities. Genetics 232, iyaf215 (2026).

35. wwPDB consortium. Protein Data Bank: the single global archive for 3D macromolecular structure data. Nucleic Acids Res. 47, D520–D528 (2019).

36. Arafeh, R., Shibue, T., Dempster, J. M., Hahn, W. C. & Vazquez, F. The present and future of the Cancer Dependency Map. Nat. Rev. Cancer 25, 59–73 (2025).

37. Varadi, M. et al. 3D-Beacons: decreasing the gap between protein sequences and structures through a federated network of protein structure data resources. GigaScience 11, giac118 (2022).

38. Magaña, P., Nair, S., Varadi, M. & Velankar, S. Harnessing the 3D-Beacons Network: A Comprehensive Guide to Accessing and Displaying Protein Structure Data. Curr. Protoc. 4, e1047 (2024).

39. Abramson, J. et al. Accurate structure prediction of biomolecular interactions with AlphaFold 3. Nature 630, 493–500 (2024).

40. Dingus, J. G., Tang, J. C., Amamoto, R., Wallick, G. K. & Cepko, C. L. A general approach for stabilizing nanobodies for intracellular expression. eLife 11, e68253 (2022).

41. Cock, P. J. A. et al. Biopython: freely available Python tools for computational molecular biology and bioinformatics. Bioinformatics 25, 1422–1423 (2009).

42. Kabsch, W. & Sander, C. Dictionary of protein secondary structure: Pattern recognition of hydrogen-bonded and geometrical features. Biopolymers 22, 2577–2637 (1983).

43. Logan, J., Sumner, J., Grigas, A., Shattuck, M. & OHern, C. The Effect of Stereochemical Constraints on the Radius of Gyration of Folded Proteins. (2025). doi:10.48550/arXiv.2501.02424.

44. Jumper, J. et al. Highly accurate protein structure prediction with AlphaFold. Nature 596, 583–589 (2021).

45. Dauparas, J. et al. Robust deep learning–based protein sequence design using ProteinMPNN. Science 378, 49–56 (2022).

46. Goverde, C. A. et al. Computational design of soluble and functional membrane protein analogues. Nature 631, 449–458 (2024).

47. Feuermann, M. et al. A compendium of human gene functions derived from evolutionary modelling. Nature 640, 146–154 (2025).

48. Biocompare. https://www.biocompare.com/Antibodies/.

49. Velankar, S. et al. SIFTS: Structure Integration with Function, Taxonomy and Sequences resource. Nucleic Acids Res. 41, D483–D489 (2013).

50. Dana, J. M. et al. SIFTS: updated Structure Integration with Function, Taxonomy and Sequences resource allows 40-fold increase in coverage of structure-based annotations for proteins. Nucleic Acids Res. 47, D482–D489 (2019).

51. Lee, B. & Richards, F. M. The interpretation of protein structures: estimation of static accessibility. J. Mol. Biol. 55, 379–400 (1971).

52. Mitternacht, S. FreeSASA: An open source C library for solvent accessible surface area calculations. F1000Research 5, 189 (2016).

53. Steinegger, M. et al. HH-suite3 for fast remote homology detection and deep protein annotation. BMC Bioinformatics 20, 473 (2019).

54. Pupko, T., Bell, R. E., Mayrose, I., Glaser, F. & Ben-Tal, N. Rate4Site: an algorithmic tool for the identification of functional regions in proteins by surface mapping of evolutionary determinants within their homologues. Bioinformatics 18 Suppl 1, S71–77 (2002).

55. Pegerto. pegerto/pyrate4site. https://github.com/pegerto/pyrate4site (2025).

56. Shrake, A. & Rupley, J. A. Environment and exposure to solvent of protein atoms. Lysozyme and insulin. J. Mol. Biol. 79, 351–371 (1973).

57. Tien, M. Z., Meyer, A. G., Sydykova, D. K., Spielman, S. J. & Wilke, C. O. Maximum Allowed Solvent Accessibilites of Residues in Proteins. PLOS ONE 8, e80635 (2013).

58. Zhang, C., Shine, M., Pyle, A. M. & Zhang, Y. US-align: universal structure alignments of proteins, nucleic acids, and macromolecular complexes. Nat. Methods 19, 1109–1115 (2022).

